# Structure of Factor VII Gla domain bound to EPCR

**DOI:** 10.64898/2025.12.18.695140

**Authors:** Jacinto López-Sagaseta, María Gilda Dichiara-Rodríguez

## Abstract

The precise molecular determinants that underlie the binding of activated factor VII (FVIIa) to the endothelial cell protein C receptor (EPCR) have long remained unresolved. We report here the crystal structure of factor VII Gla domain bound to human EPCR, revealing a binding geometry in essence indistinguishable from that of the PC-EPCR interface and confirming absolute structural competition for EPCR binding. Relative to previous FVII structures, we observe receptor-dependent repositioning of the Gla ω loop coupled to a selective arrangement in which metal ion at position four adopts a Ca^2+^-compatible and EPCR-restricted coordination state. These findings resolve a long-standing gap and inform the structural basis for the binding of EPCR to FVIIa, a current therapeutic intervention in hemophilia.

## Introduction

Over the past few decades, many groups have explored a potential interaction of EPCR with other Gla-containing factor, yet the debate gained clinical relevance with activated factor VII (FVIIa)^1–5^. This attention arose from the high circulating levels achieved upon administration of recombinant FVIIa (rFVIIa)^6^, a current pharmacological intervention in the treatment of hemophilia. This scenario raised the question of whether the EPCR-FVIIa interaction could play an important role in the hemostatic potential of rFVIIa. *In vitro*, EPCR can inhibit tissue factor-dependent FVIIa coagulant activity^1^. *In vivo*, it was found that EPCR potentiates rFVIIa’s hemostatic effect by competing with protein C^4^.

Despite numerous studies have addressed the EPCR-FVIIa interaction, including mutational studies at the EPCR-FVII interface^1^, a structure that confirms a coherent EPCR-FVIIa docking and provides the binding signature had not yet been reported and remained elusive. Here, we present the X-ray structure of FVII/FVIIa Gla domain bound to human EPCR, revealing the footprint of the interaction and a side-by-side comparison with that of the established EPCR-PC/APC interface.

## Results and discussion

The crystal structure of EPCR with bound FVII Gla domain was solved at 3.0 Å resolution, with four EPCR-FVII Gla complexes in the asymmetric unit (Fig. 1 and Supplementary Table 1). Unambiguous electron density enabled the identification of the docking mode and the binding interface, including the ω-loop and Ca^2+^ ions that are part of FVII Gla domain. EPCR presents its canonical CD1-like fold, including the hydrophobic tunnel filled with a phospholipid (Fig. 1A). The bound FVII Gla is located at one of receptor’s ends, and binds EPCR through ionic and H-bonds mediated by Gla7, Gla25 and Gla29, and indirectly, via Ca^2+^ coordination (EPCR Glu86 with Gla 7, 26 and 29). Essentially replicating PC docking geometry (Fig. 1B), FVII Gla ω-loop contributes with non-polar interactions, triggering a conformational change in EPCR Arg156, that releases space for accommodation of Phe4, tightly positioned adjacent to EPCR Gln75 and Tyr154 (Figs. 1A and 1C). Leu5 and Leu8 are also located at a distance suitable for participating with hydrophobic and Van der Waals contacts. EPCR’s hydrophobic groove houses a bound phospholipid^7–10^, whose phosphate group, structurally exposed outwardly, attracts Arg156 side chain through electrostatic interactions (Fig. 1A). Overall, the binding mode is indistinguishable from that previously observed for protein C Gla domain (Fig. 1B). Likewise, a set of up to seven divalent cations, mainly Ca^2+^ ions, are distributed within FVII Gla residues, coordinating and driving a stable conformation that provides, ultimately, a productive binding to EPCR. The presence of Mg^2+^ ions had been previously reported to enhance the binding to EPCR^11,12^, therefore, MgCl_2_ was included in the crystallization conditions on purpose. We observed strong Fo-Fc electron density for five ion binding sites consistent with a scattering power for Ca^2+^ ions rather than Mg^2+^ (Fig. 1D). Two additional sites with weaker Fo-Fc peaks were more ambiguous as they could represent Mg^2+^ with full occupancy, but also Ca^2+^ with partial occupancy (Fig. 1D). These results agree with previous reports that demonstrated the binding of both types of ions to PC and FVII Gla^7,12,13^.

**Fig. 1.**
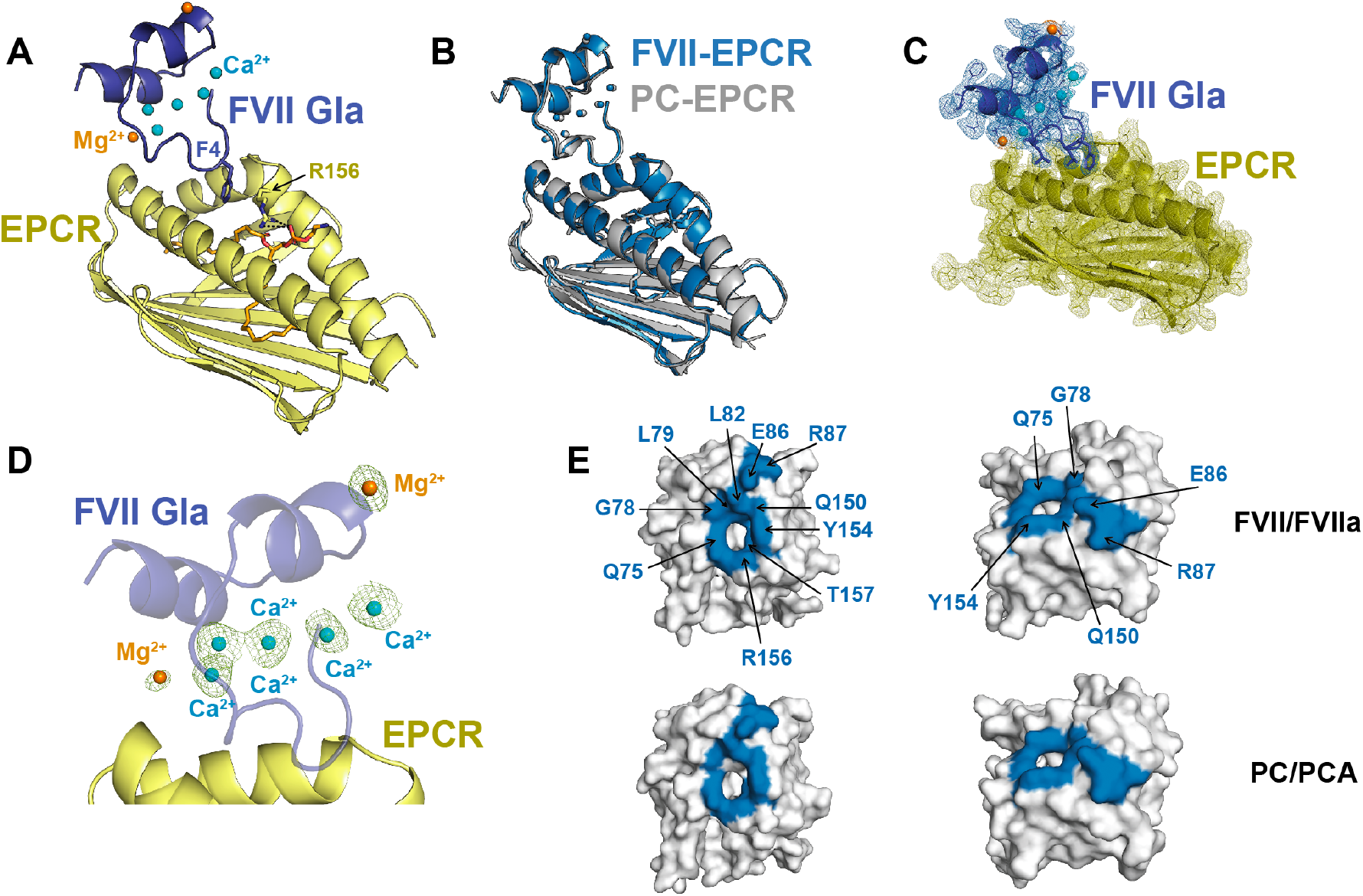
Structural basis for FVII Gla domain binding to EPCR. **(A)** FVII Gla domain (light blue) is shown bound to EPCR (yellow), with coordinating Ca^2+^ and Mg^2+^ ions displayed as cyan and orange spheres, respectively. N-glycosylation at position Asn119 in EPCR is represented as sticks. The phospholipid molecule, also shown in sticks, is bound inside the groove formed by the two EPCR alpha helices. (B) Structural superposition of the FVII Gla– EPCR complex (blue) with that of the PC Gla-EPCR complex, PDB 1LQV (gray). (C) The FVII Gla domain–EPCR structure is displayed in combination with experimental electron density. The 2Fo–Fc electron density maps are contoured at 1.0 σ (blue mesh for FVII Gla, yellow for EPCR), illustrating the experimentally-determined docking and well-defined signal for both the FVII Gla domain and EPCR molecules at the binding interface. (D) Positive Fo-Fc electron density (green) determined by omitting the Ca^2+^ and Mg^2+^ ions (cyan and orange spheres, respectively) that contribute to FVII Gla overall fold and EPCR binding. (E) Two views of the residues in EPCR (shown as a grey surface) that directly participate in contacts with FVII Gla are indicated and highlighted in blue color. For reference, the footprint of PC Gla is also shown.

Structural alignment of FVII Gla-EPCR and PC Gla-EPCR (PDB 1LQV) complexes with EPCR as reference frame reveals a highly accurate superposition (0.305 Å) of both Gla domains. The ω-loops in each Gla shows almost perfect overlap and orientation relative to EPCR, as well as near identical footprints on the receptor’s surface (Fig. 1E). The overall docking geometry presents a buried surface area of 468 Å^2^ for the binding interface (Fig. 1E), which, again, is similar with that observed in the PC Gla-EPCR complex (467 Å^2^), suggesting that FVII Gla-EPCR interaction is solid and energetically coherent. An additional contact was observed between Gln85 and the backbone oxygen of Gla7 in the FVII Gla–EPCR complex. However, this interaction occurs at a distance of ∼ 4 Å and is present in only one of the four complexes within the asymmetric unit; therefore, it is unlikely to represent a relevant interaction.

Prior alanine-scan mutagenesis^1^ found a limited number of residues in EPCR critical for FVIIa binding, with many being equivalent to those identified structurally as key for the PC Gla-EPCR interaction^7^. This suggested that FVII/FVII Gla engagement on EPCR shared with PC a highly similar binding site on EPCR’s surface. The FVII Gla-EPCR structure shows that Leu82, Glu86, and Arg87, all located in EPCR a1 helix, conform a cluster of direct contacts with the FVII Gla. Leu82 participates with hydrophobic packing interactions with FVII Gla7 and Leu8 backbone atoms, whereas Glu86 and Arg87 establish, respectively, H-bonds with Gla7 and Gla29 and salt bridges with Gla25 (Supplementary Fig. 1A). On EPCR a2 helix, Gln150 and Tyr154 establish direct H-bonds with Gla7 (Supplementary Fig. 1B), with Tyr154 also reported to be fundamental for binding of FVIIa^1^. Thus, these contacts further contribute to the overall strength of the FVII Gla-EPCR interaction. Additional contacts mediated by a coordinating Ca^2+^ ion were observed, which links Glu86 in EPCR with Gla26 in FVII Gla domain, and creates a more robust network of interactions with Gla7 and Gla29, in all cases, in a manner highly similar with that of the PC-EPCR interface (Supplementary Fig. 1C). Phe146 and Arg158, also pivotal for FVII binding^1^, contribute internally to a stable EPCR architecture. Phe146, close to EPCR Leu82 and Val83, plays a stabilizing role in a local hydrophobic site underneath FVII Gla binding surface, while Arg158’s side chain engages Arg130 and Ala131 backbone atoms in an adjacent EPCR β-strand, thereby supporting the structural integrity of EPCR.

Like PC, the EPCR-FVIIa interface is also dominated by non-polar intermolecular contacts. Our structure reveals precise interactions between EPCR and the FVIIa Gla domain (Supplementary Fig. 2). Electron density supports docking of the Gla w loop onto the EPCR surface adjacent to the bound lipid, with Phe4, Leu5, and Leu8 located at the core of the binding interface (Supplementary Fig. 2A). Surface comparisons show that FVII Gla occupies a similar EPCR binding pocket as protein C, adopting an analogous docking pose (Supplementary Fig. 2B). We also observed conserved hydrophobic and Van der Waals contacts at the binding interface. These involve Gln75 and Arg156 in EPCR and are mediated by the Gla w loop Phe4 and Leu5 residues, as well as Leu8, beyond this loop (Supplementary Fig. 2C). A particular attention is deserved to Tyr154, which provides a dense set of non-polar interactions with the Gla N-terminal region (Supplementary Fig. 2D).

A structural aligment of the EPCR-bound and EPCR-free (PDB ID 2A2Q) FVII Gla domains reveal molecular plasticity at the N-terminal ω-loop region (Fig. 2A). It is observed that EPCR engagement is restricted to a repositioning of residues Phe4, Leu5, and Leu8 to enable a distinct orientation of the ω-loop towards the receptor surface. The alternative conformation takes place without global rearrangement of the Gla fold and is spatially coupled to central divalent cation coordination. The observed structural discrepancies indicate that EPCR binding is associated with selective remodeling of the N-terminal segment of the FVII Gla domain.

**Fig. 2.**
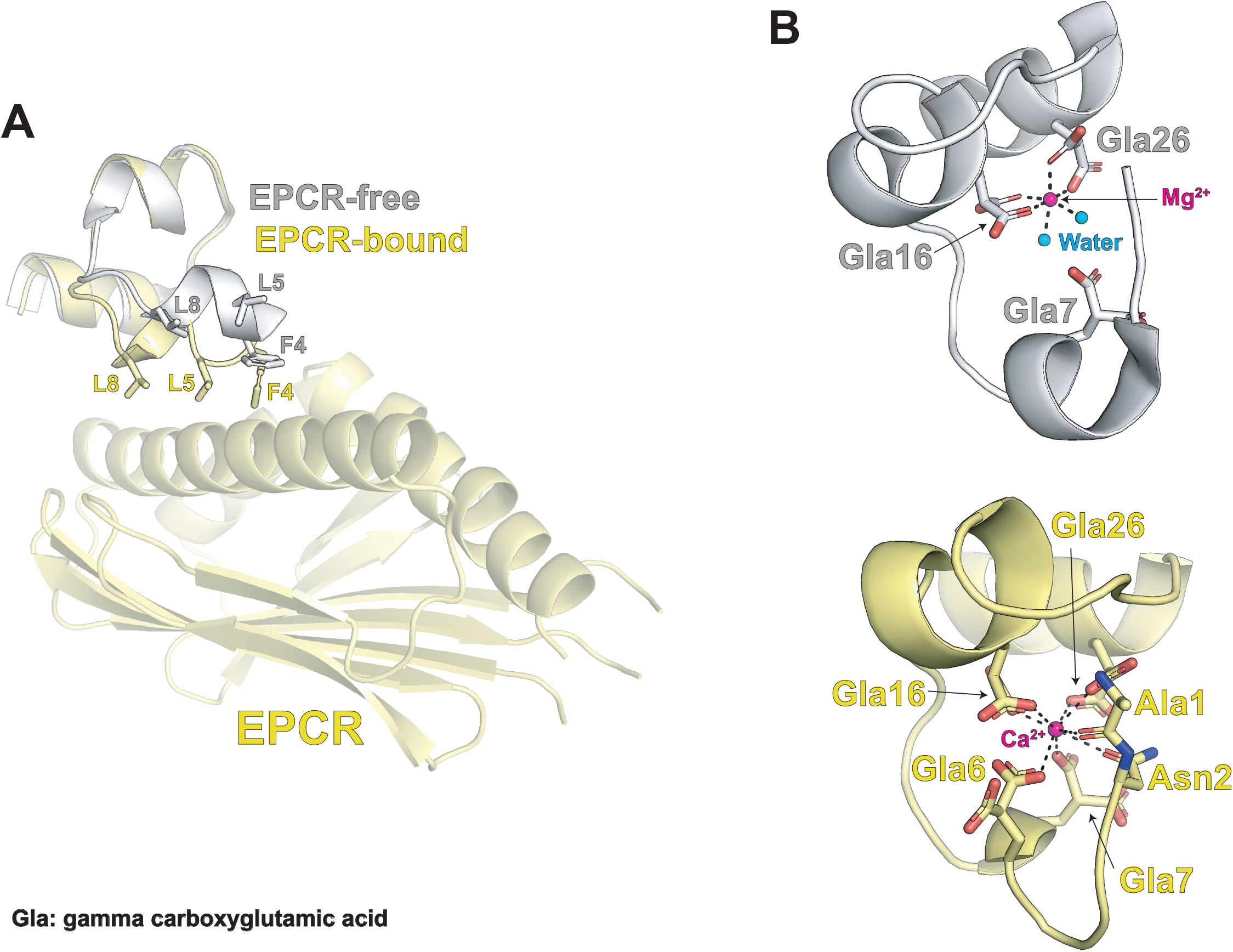
EPCR binding determines FVIIa Gla conformation in a fine-tuned divalent cation manner. **(A)** Structural alignment of the FVII Gla domain in the EPCR-free state (gray, PDB 2A2Q) and EPCR-bound state (yellow, PDB 9TJX) highlighting conformational differences at the ω-loop. The EPCR platform is shown in pale yellow for reference. **(B)** Close-up viewillustrating the EPCR-dependent arrangement of the metal coordination environment at position 4. Coordinating interactions are indicated by dashed lines.

A previous study^14^ reported that Mg^2+^ shows a maximum displacement capacity of up to three out of the seven Ca^2+^ sites previously found in the Gla domain of FVIIa^15^. However, the 4-Ca^2+^/3-Mg^2+^ configuration is not observed in the EPCR-bound state. Structurally and chemically, the coordination geometry is most consistent with Ca^2+^ occupancy at position four, which yields a metal ion arrangement similar with that found for the EPCR-protein C interface, that is, five Ca^2+^ and two Mg^2+^ ions^12^. This Ca^2+^/Mg^2+^ configuration is supported by additional coordination for Ca^2+^ at position 4 involving Gla7, which adopts a substantially different conformation (Fig. 2B) relative to EPCR-free structures. The presence of Ca^2+^ at position four is further supported by improved R factor and Rfree values when position four is filled with Ca^2+^, consistent B-factors, and metal-oxygen distances characteristic of Ca^2+^ rather than Mg^2+^ coordination. Taken together, these findings indicate that FVIIa/Mg^2+^/Ca^2+^ complexes with different stoichiometries can coexist in solution, but only those in which site four of the Gla domain is occupied by Ca^2+^ are likely to sustain EPCR binding. This is therapeutically relevant, since recombinant FVIIa is formulated in CaCl_2_. Given that both tissue factor and EPCR interactions are stabilized by the presence of Mg^2+^, these findings raise the possibility that targeted modulation of the Gla-domain metal coordination could potentiate receptor engagement.

The FVII Gla-EPCR crystal structure provides definitive evidence for a genuine FVII-EPCR interaction and addresses a long-lasting question pertaining EPCR restriction to PC/APC. Our crystal structure confirms that EPCR is not restricted to PC/APC, but instead, can be targeted by the Gla domain of FVII through a highly conserved docking mode. Our data further suggest a diverse population of FVIIa-Ca^2+^/Mg^2+^ assemblies in solution, with EPCR binding being favored by those where site four in the Gla domain is filled with Ca^2+^, highlighting fine-tuned metal-driven modulation of FVIIa behaviour.

Pharmacological interventions with rFVIIa lead to a notable elevation in plasma levels, boosting the potential for EPCR to capture FVIIa instead of PC. In agreement with previous reports, this molecular competition would have an impact in the generation rate of APC, and, consequently, in the anticoagulant system.

## Methods

### Protein expression

The EPCR ectodomain (N30Q to reduce N-glycosylation) was cloned into the pcDNA3.4 secretion vector downstream an N-terminal TwinStrepTag® followed by a 3C protease site. EPCR was isolated from the cell culture supernatant upon transient-expression in CHO-S cells, with a Strep-Tactin affinity column followed by size-exclusion chromatography. Tag-free and deglycosylated EPCR was achieved by incubation with 3C protease and PNGase (produced in-house), with the enzymes being removed through a NiNTA resin prior to concentration and storage of EPCR target protein.

### Crystallization of FVII Gla-EPCR complex

Pure, tag-free and deglycosylated EPCR was mixed with a 3-fold molar excess of FVII Gla peptide in the presence of CaCl_2_ and MgCl_2_. The protein-peptide complex was recovered via size-exclusion chromatography and concentrated prior to crystallization. Crystals of FVII Gla-EPCR complex were obtained by the sitting-drop vapor diffusion at room temperature and cryoprotected with glycerol before freezing.

### Resolution of the FVII Gla-EPCR complex structure

X-ray diffraction data were collected at Xaira beamline. Diffraction spots were processed with XDS^16^ and Aimless^17^ packages. Structure solution was achieved via molecular replacement with Phaser^18^, using EPCR (PDB 1L8J) and FVII Gla domain (1DAN) atomic coordinates as reference models. The structure was refined with Phenix.refine^19^, with iterative manual building in Coot^20^.

## Acknowledgements

We are grateful to the staff of Xaira beamline at ALBA Synchrotron for their assistance with X-ray diffraction data collection.

## Autorship contributions

Conceived research: JLS; Performed experiments: JLS, GDR; Data collection and analysis: JLS; Draft writing: JLS.

## Data availability

Atomic coordinates and structure factors have been deposited and are available at the Protein Data Bank under the accession code PDB ID 9TJX.

## References

1. Lopez-Sagaseta, J. et al. Binding of factor VIIa to the endothelial cell protein C receptor reduces its coagulant activity. J Thromb Haemost 5, 1817–1824 (2007).

2. Preston, R. J. et al. Multifunctional specificity of the protein C/activated protein C Gla domain. J Biol Chem 281, 28850–28857 (2006).

3. Ghosh, S., Pendurthi, U. R., Steinoe, A., Esmon, C. T. & Rao, L. V. Endothelial cell protein C receptor acts as a cellular receptor for factor VIIa on endothelium. J Biol Chem 282, 11849–11857 (2007).

4. Keshava, S. et al. Factor VIIa interaction with EPCR modulates the hemostatic effect of rFVIIa in hemophilia therapy: Mode of its action. Blood Adv 1, 1206–1214 (2017).

5. Disse, J. et al. The endothelial protein C receptor supports tissue factor ternary coagulation initiation complex signaling through protease-activated receptors. J Biol Chem 286, 5756–5767 (2011).

6. Roberts, H. R., Monroe, D. M. & White, G. C. The use of recombinant factor VIIa in the treatment of bleeding disorders. Blood vol. 104 3858–3864 Preprint at 10.1182/blood-2004-06-2223 (2004).

7. Oganesyan, V. et al. The crystal structure of the endothelial protein C receptor and a bound phospholipid. J Biol Chem 277, 24851–24854 (2002).

8. Lopez-Sagaseta, J. et al. sPLA2-V inhibits EPCR anticoagulant and antiapoptotic properties by accommodating lysophosphatidylcholine or PAF in the hydrophobic groove. Blood 119, 2914–2921 (2012).

9. Erausquin, E. et al. Identification of a broad lipid repertoire associated to the endothelial cell protein C receptor (EPCR). Sci Rep 12, (2022).

10. Erausquin, E. et al. Structural vulnerability in EPCR suggests functional modulation. Sci Rep 14, (2024)

11. López-Sagaseta, J., Montes, R. & Hermida, J. Recombinant expression of biologically active murine soluble EPCR. Protein Expr Purif 64, 194–197 (2009).

12. Vadivel, K. et al. Structural and functional studies of γ-carboxyglutamic acid domains of factor VIIa and activated protein C: Role of magnesium at physiological calcium. J Mol Biol 425, 1961–1981 (2013).

13. Lopez-Sagaseta, J., Montes, R. & Hermida, J. Recombinant expression of biologically active murine soluble EPCR. Protein Expr Purif 64, 194–197 (2009).

14. Bajaj, S. P., Schmidt, A. E., Agah, S., Bajaj, M. S. & Padmanabhan, K. High resolution structures of p-aminobenzamidine- and benzamidine-VIIa/soluble tissue factor: Unpredicted conformation of the 192-193 peptide bond and mapping of Ca2+, Mg2+, Na+, and Zn2+ sites in factor VIIa. Journal of Biological Chemistry 281, 24873–24888 (2006).

15. D W Banner, A. D. C. C. F. K. W. A. G. W. H. K. Y. N. D. K. The crystal structure of the complex of blood coagulation factor VIIa with soluble tissue factor. Nature 380, 41–6 (1996).

16. Kabsch, W. XDS. Acta Crystallogr D Biol Crystallogr 66, 125–132 (2010).

17. Evans, P. R. & Murshudov, G. N. How good are my data and what is the resolution? Acta Crystallogr D Biol Crystallogr 69, 1204–1214 (2013).

18. McCoy, A. J. et al. Phaser crystallographic software. J Appl Crystallogr 40, 658–674 (2007).

19. Adams, P. D. et al. PHENIX: a comprehensive Python-based system for macromolecular structure solution. Acta Crystallogr D Biol Crystallogr 66, 213–221 (2010).

20. Emsley, P., Lohkamp, B., Scott, W. G. & Cowtan, K. Features and development of Coot. Acta Crystallogr D Biol Crystallogr 66, 486–501 (2010).

