## Supplementary materials for "Structure of Factor VII Gla domain bound to EPCR"

### SUPPLEMENTARY MATERIAL

**METHODS**

**SUPPLEMENTARY FIGURE LEGENDS**

**SUPPLEMENTARY TABLE 1. DIFFRACTION DATA COLLECTION AND REFINEMENT STATISTICS.**

#### METHODS (EXTENDED)

*Production and purification of EPCR*

The extracellular region of EPCR was PCR amplified from a previous EPCR construct carrying the N30Q mutation and designed to obtain a reduced degree of N-glycosylation. Oligos were designed to introduce pspOMI (5`) and HindIII (3`) sites. The amplified product was purified from agarose gel, digested with the restriction enzymes (Fisher Scientific) and used for ligation with T4 DNA ligase (Thermo Fisher Scientific) into an in-house version of pcDNA3.4 containing a secretion signal, a TwinStrepTag and a 3C site upstream EPCR sequence. Competent *E. coli* DH5α cells were transformed with the ligation product, positive transformants were identified and grown in 10 mL LB supplemented with ampicillin (50 μg/mL, Fisher). The recombinant plasmid was isolated with the GeneJET Plasmid Miniprep Kit (Thermo Fisher Scientific) following the manufacturer's instructions, and the target sequence was verified via Sanger sequencing. Transfection-grade DNA was prepared from 100 mL of LB culture supplemented with ampicillin (50 μg/mL, Fisher) using the PureLink$^{TM}$ HiPure Plasmid Midiprep Kit (Thermo Fisher) according to the manufacturer's instructions.

CHO-S cell (Invitrogen) were cultured in ExpiCHO$^{TM}$ Expression Medium (Thermo Fisher) and transfected with the recombinant EPCR plasmid using the ExpiFectamine$^{TM}$ CHO Transfection Kit (Thermo Fisher). Transfected cells were then incubated with orbital agitation at 37ºC and 8% $CO_2$. Eight days post-transfection, the cell supernatant was freed of cells by centrifugation (10000xg, 20 min), filtered through a 0.45 μm filter (Merck) to eliminate any particulate contaminants and buffer exchanged to 100 mM Hepes pH 7.4, 150 mM NaCl (HBS), 1 mM EDTA in a HiPrep 26/10 column (Cytiva) connected to an AKTA Pure System (Cytiva). The resulting sample was then loaded onto a 1-ml Strep-Tactin XT 4Flow (Fisher Scientific) column previously equilibrated in 100 mM Hepes pH 7.4, 150 mM NaCl (HBS), 1 mM EDTA. Following sample load, the column was washed with 10 column volumes and EPCR was eluted with the same buffer supplemented with 50 mM biotin. The protein was further purified via size exclusion chromatography using TBS (20 mM Tris pH 7.4, 150 mM NaCl) as mobile phase and a Superdex 200 10/300 GL column (Cytiva) column.

*Production and purification of 3C and PNGase*

The preparation of 3C protease was carried out as described earlier. PNGase was recombinantly expressed in BL21DE3 cells. DNA encoding PNGase sequence with a C-terminal 6×His tag was purchased from IDT. Both the construct and pet28 vector were double-digested with XhoI and NcoI (Fisher Scientific) following the manufacturer recommendations. Following digestion, insert and plasmids were gel-purified and ligated with T4 DNA ligase (Thermo Fisher Scientific) to generate the final plasmid. Protein expression was induced in BL21DE3 with 1 mM IPTG for 4 h and the target protein was recovered from inclusion bodies and stored frozen following previously described procedures (ref). PNGase was refolded by diluting 200 mg of inclusion bodies in 200 mL of buffer containing 2.5 M urea, 0.4 M L-arginine, 50 mM Tris pH 8, 0.5 mM oxidized glutathione, and 5 mM reduced glutathione. The sample was left stirring for 4 h at 4°C, then dyalized against 2 liters of 1X HBS pH 7.4. The surrounding buffer was replaced with fresh buffer for three times every 12 hours. The temperature was kept at 4 ºC during the whole procedure. PNGase was captured through immobilized metal affinity chromatography using a Ni-NTA resin (Teknovas), followed by size-exclusion chromatography on a Superdex 75 (S75) column equilibrated with 1X TBS pH 7.4. Pure PNGase was aliquoted, flash-frozen in liquid nitrogen and stored frozen at -80ºC.

*Removal of EPCR N-glycosylation*

The EPCR-containing fractions were pooled and subjected to a double simultaneous digestion with 3C protease and PNGase, with both enzymes being recombinanty produced and purified in our laboratory as described above for PNGase, an earlier for 3C protease (ref). EPCR was incubated with PNGase and 3C at an enzyme:substrate ratio of 1:10 and 1:100 (weight/weight), respectively, for 18 hours at 4º C. The EPCR product showed enhanced migration in SDS-PAGE (not shown), confirming complete 3C digestion and partial N-deglycosylation. The final EPCR sample was freed of enzymes by loading the samples onto a Ni-NTa resin (Teknovas). The flowthrough containing the EPCR protein was concentrated with a 10-MWCO Amicon concentrator (Millipore) and preserved frozen at -80ºC.

Preparation of the FVII Gla domain-EPCR complex

Pure, tag-free and deglycosylated EPCR was mixed with a three-fold molar excess of FVII Gla peptide (custom-synthesized, QYAOBIO) reconstituted in 20 mM Tris pH 7.4, 150 mM NaCl, 10 mM $CaCl_2$ and 6 mM $MgCl_2$. The sample was incubated for 1 hour at room temperature, then injected onto a S75 10 300 column (Cytiva) to remove the excess of peptide. The protein-peptide complex was concentrated with a 10-MWCO Amicon concentrator (Millipore) to 5.4 mg/ml. Protein concentration was determined by absorbance at 280 nm, using an extinction coefficient determined from the full protein-peptide sequence by Expasy ProtParam tool.

*Crystallization of the EPCR:FVII Gla complex*

Protein crystallization was accomplished using the sitting-drop vapor diffusion method at RT. A Crystal Gryphon LCP platform (Dunn Labortechnik GmbH) robot assisted with drop setup in 96-well crystallization plates. Drops were composed of 300 nL of protein sample and 300 nL of crystallization reagent. The crystallization plates were incubated at RT and the drops were inspected for crystal growth. Thin, clustered and plate-looking crystals appeared after three weeks in 0.2 M ammonium sulfate, 0.1 M Tris PH 8.5, 12 % w/v PEG 8000. Individual crystals were transferred to fresh drops with equal composition and additional 20 % (V/V) glycerol. After soaking, individual crystals were cryo-cooled in liquid nitrogen prior to diffraction analysis.

*Diffraction analysis, data processing and structure determination and refinement*

X-ray data was collected at Xaira beamline (Cerdanyola del Vallès, Barcelona, Spain). Diffraction spots were indexed with XDS, then scaled and merged with Aimless. The resolution cutoff was set to 3.0 Å, based on an overall completeness of 93%, corresponding to $I/\sigma(I)$ and CC1/2 values of 2.2 and of 0.705, respectively, in the highest-resolution shell. The EPCR:FVII Gla structure was solved via molecular replacement using Phaser using, on the one hand, the EPCR atomic coordinates from PDB 1L8J, omitting the phos and any water or small molecule. On the other hand, the atomic coordinates of the FVII Gla domain from PDB 1DAN, with ions omitted. Solution was achieved with a combined search of four EPCR molecules and three Gla domain molecules. The fourth Gla domain molecule was manually added. Refinement was performed using Phenix.refine with the non-crystallographic symmetry (NCS) parameter enabled. A series of manual model building in Coot, guided by Fo-Fc and 2Fo-Fc electron density maps, and refinement cycles were undertaken to generate the final structure.

**Supplementary Figure 1. Comparison of Gla domain-EPCR polar contacts in PC and FVII complexes.** (A) H-bonds bonds and salt bridges (black dashed lines) established between Glu86 and Arg87 in EPCR with several Gla residues in FVII and PC Gla domains, including Gla7, Gla9 and Gla25. (B) H-bonds mediated by Gla7 residues in PC (gray) and FVII (purple) complexes are indicated. All residues that participate in the interactions are indicated and highlighted as sticks. (C) Additional $Ca^{2+}$-dependent contacts formed by Glu86 with Gla7 and Gla29 but also with Gla26. $Ca^{2+}$ ions are shown in gray (PC complex) and blue color (FVII complex). The PC complex highlights also a water molecule (cyan) coordinated by the $Ca^{2+}$ ion.

**Supplementary Figure 2. Conserved hydrophobic interactions are critical for FVII-EPCR binding.** (A) Close-up view of the FVII Gla N-terminal ω loop bound to EPCR. The FVII Gla backbone is displayed as sticks in purple and EPCR as cartoon in yellow. The gray mesh indicates 2Fo–Fc electron density, while the green mesh represents positive Fo–Fc difference density, confirming the location of FVII ω loop Phe4 in between EPCR alpha helices. (B) Surface representation of EPCR in yellow color, with hydrophobic residues (Ala/Val/Leu/Ile/Met/Phe/Tyr/Trp) highlighted in gray. The PC and FVII Gla domains are shown in cartoon mode bound to EPCR with the same color configuration. Side-by-side comparison of Phe4, Leu5 and Leu8 residues of the PC and FVII ω loops, highlighted as sticks and occupying a hydrophobic pocket in EPCR upper region. (C) Non-polar contacts (dashed lines) established between EPCR and the Gla domains of PC (gray) and FVII (blue). (D) Tyr154 mediates a dense set of Van der Waals and hydrophobic interactions with the Gla domain N-terminal residues in both PC and FVII complexes, indicating the highly conserved binding mechanism.

**Supplementary Table 1 |** Diffraction data collection and refinement statistics.

| DATA COLLECTION | FVII GLA-EPCR COMPLEX |
|---|---|
| Resolution range (Å) | 43.67 - 3.00 (3.18 - 3.00) |
| Space group | C121 |
| Unit cell | 278.41 44.22 111.25 90 93.01 90 |
| Total reflections | 77200 (13691) |
| Unique reflections | 25876 (4440) |
| Multiplicity | 3.0 (3.1) |
| Completeness (%) | 93.0 (99.6) |
| Mean I/sigma(I) | 4.2 (2.2) |
| Wilson B-factor | 48.83 |
| R-merge | 0.153 (0.480) |
| R-meas | 0.187 (0.583) |
| R-pim | 0.106 (0.326) |
| CC1/2 | 0.967 (0.705) |
| Reflections in refinement | 25875 (3038) |
| Reflections (R-free) | 1249 (148) |
| R-work | 0.239 (0.254) |
| R-free | 0.290 (0.316) |
| RMS(bonds) | 0.002 |
| RMS(angles) | 0.66 |
| Ramachandran favored (%) | 98.98 |
| Ramachandran allowed (%) | 0.87 |
| Ramachandran outliers (%) | 0.15 |
| Average B-factor ($Å^2$) | 54.61 |

Statistics for the highest-resolution shell are shown in parentheses.
