## Supplementary figures and images for "Structure of Factor VII Gla domain bound to EPCR"

### Supplementary Figure 1

#### Supplementary Figure 1

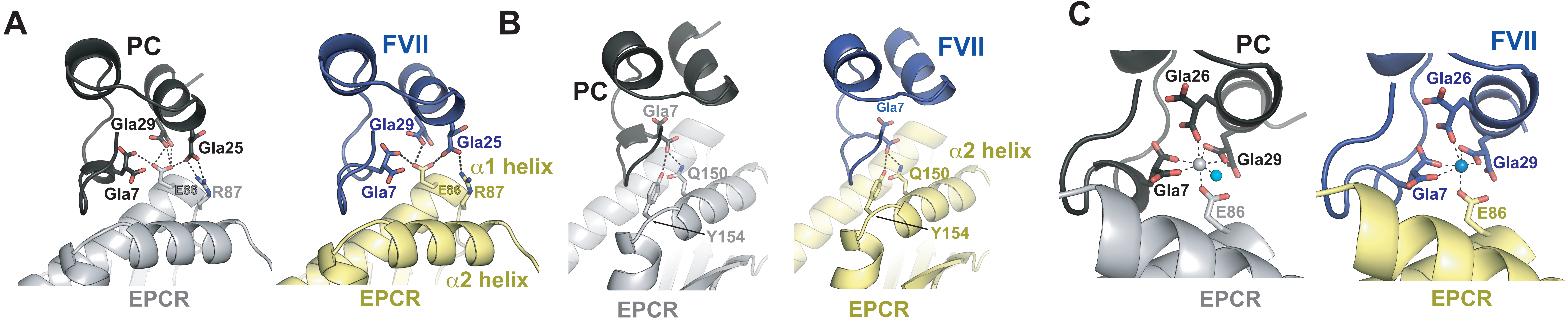

### Supplementary Figure 2

#### Supplementary Figure 2

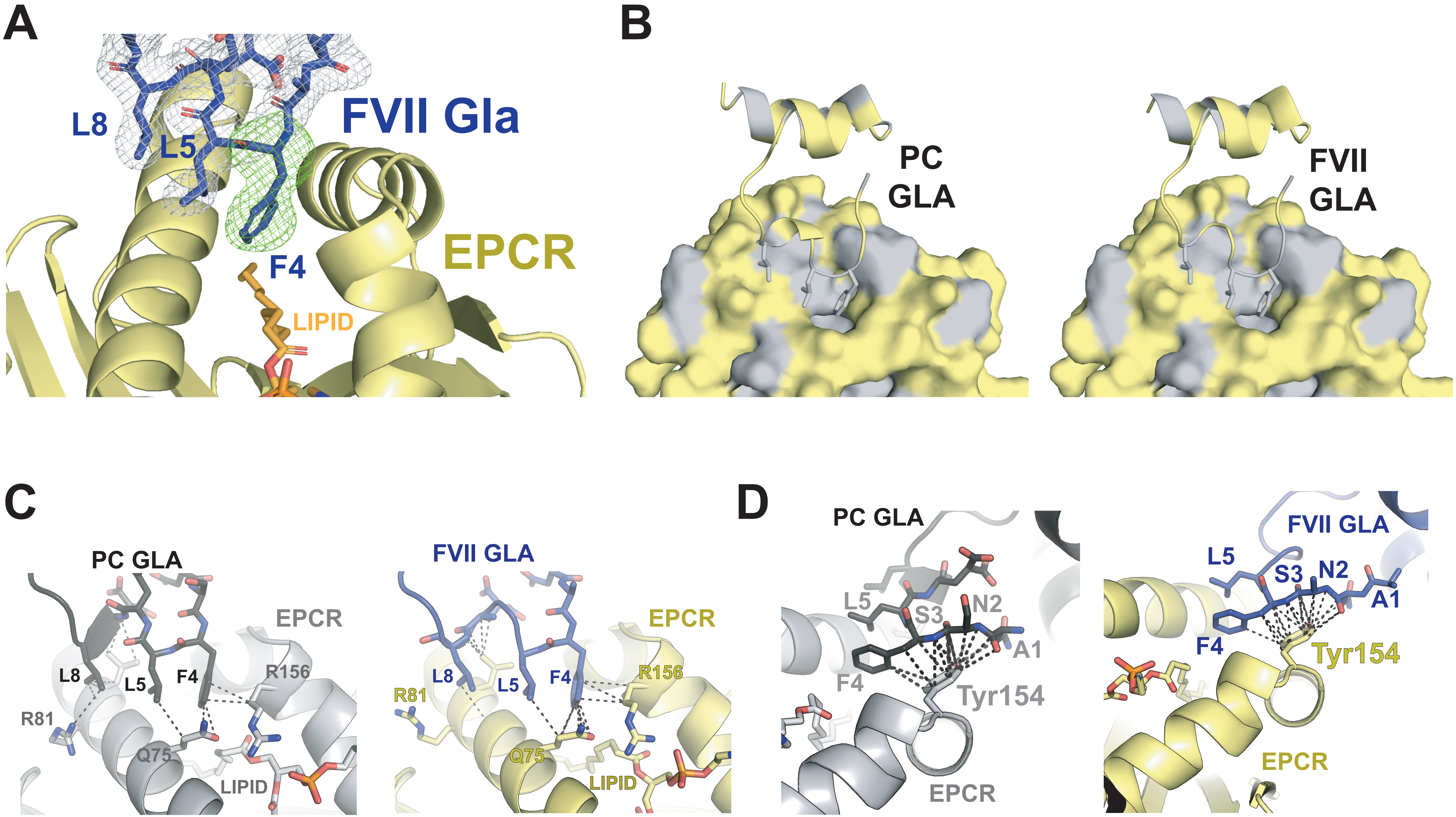
